# A comparison of six DNA extraction protocols for 16S, ITS, and shotgun metagenomic sequencing of microbial communities

**DOI:** 10.1101/2022.03.07.483343

**Authors:** Justin P. Shaffer, Carolina S. Carpenter, Cameron Martino, Rodolfo A. Salido, Jeremiah J. Minich, MacKenzie Bryant, Karenina Sanders, Tara Schwartz, Gregory Humphrey, Austin D. Swafford, Rob Knight

## Abstract

Microbial communities contain a broad phylogenetic diversity of organisms, however the majority of methods center on describing bacteria and archaea. Fungi are important symbionts in many ecosystems, and are potentially important members of the human microbiome, beyond those that can cause disease. To expand our analysis of microbial communities to include fungal ITS data, we compared five candidate DNA extraction kits against our standardized protocol for describing bacteria and archaea using 16S rRNA gene amplicon- and shotgun metagenomics sequencing. We present results considering a diverse panel of host-associated and environmental sample types, and comparing the cost, processing time, well-to-well contamination, DNA yield, limit of detection, and microbial community composition among protocols. Across all criteria, we found the MagMAX Microbiome kit to perform best. The PowerSoil Pro kit performed comparably, but with increased cost per sample and overall processing time. The Zymo MagBead, NucleoMag Food, and Norgen Stool kits were included.

**Accession numbers:** Raw sequence data were deposited at the European Nucleotide Archive (accession#: ERP124610) and raw and processed data are available at Qiita (Study ID: 12201). All processing and analysis code is available on GitHub (github.com/justinshaffer/Extraction_kit_testing).

**Methods summary:** To allow for downstream applications involving fungi in addition to bacteria and archaea, we compared five DNA extraction kits with our previously established, standardized protocol for extracting DNA for microbial community analysis. Across ten diverse sample types, we found one extraction kit to perform comparably or better than our standardized protocol. Our conclusion is based on per-sample comparisons of DNA yield, the number of quality-filtered sequences generated, the limit of detection of microbial cells, microbial community alpha-diversity, beta-diversity, and taxonomic composition, and extent of well-to-well contamination.

## Introduction

Research into microbial communities continues to reveal links that help to support both human and environmental sustainability [1-4]. In order to properly identify important connections between microbiota and human health and well-being [5-7], parallel work must foster the innovation of methods that refine our view of microbial communities [4, 8-10]. However, widespread adoption of standardized methods for performing microbiome studies continues to be hindered by a lack of approaches that capture information from all organisms, or from across diverse sample types [11-13].

Whereas methods for capturing information from bacteria and archaea have been well-developed and widely adopted, those that additionally consider microbial eukaryotes such as fungi have received less attention [14-16]. Similar to bacteria and viruses for mammals, fungi are the most important plant pathogens worldwide [17,18] and represent an increasing threat to certain groups of animals, including amphibians [19-21]. Fungi are also invaluable components of soils and forests [22-24], and are currently emerging as important members of the human microbiome (*i*.*e*., mycobiome) [25-27]. Currently, our protocol for DNA extraction for high-throughput microbiome sequencing focuses on describing bacterial/archaeal taxa, and has not yet been tested to additionally describe Fungi.

Here, we aimed to identify an extraction kit that extracts DNA from fungal communities while also producing DNA output and community composition for bacteria/archaea similar to our previously established, standardized protocol [28]. We compared DNA yield, the number of quality sequences, microbial community alpha- and beta-diversity and taxonomic composition, and also technical differences in the limits of detection of bacterial and fungal cells [29], and the extent of sample-to-sample (*i*.*e*., well-to-well) contamination [30-32] among extraction protocols.

## Materials & Methods

### Sample collection

To compare each candidate extraction kit against our standardized protocol, we collected a wide selection of samples from human body sites and the environment and centered on types common studies of microbial communities, following Marotz *et al*. (2017) and Shaffer *et al*. (2021) [28,33].This set of sample types and protocols for collecting each, the “Earth Microbiome Project (EMP) in a box”, was drafted for widespread use in benchmarking and similar studies [33]. For this study and following Shaffer *et al*. (2021), we included a total of 6 human skin samples, 6 human oral samples, 4 built environment samples, 10 fecal samples, 6 human urine samples, 2 human breastmilk samples, 6 soil samples, 4 water samples, 4 fermented food samples, and 2 tissue samples. Except where described otherwise below, we collected samples using Puritan wood-handled, cotton swabs, following the EMP standard protocol [14,33].

We collected samples in a way that allowed technical replication across extraction protocols (*i*.*e*., three technical replicates per protocol), and aliquoted each unique sample across all extraction kits for comparison of extraction efficiency, following Marotz *et al*. (2017) and Shaffer *et al*. (2021) [28,33]. Human skin samples included those from the foot and armpit, which were collected from three individuals by rubbing five cotton swabs simultaneously on the sole of each foot or armpit, respectively, for 30 s. Human oral samples included saliva, which was collected from 12 individuals by active spitting into a 50 mL centrifuge tube. Built environment samples included floor tiles (0.3 m^2^) from each of two separate laboratory bays, which were sampled separately with nine cotton swabs rubbed simultaneously across one tile surface for 30 s, and computer keyboards from each of two individuals also sampled with nine cotton swabs for 30 s. Fecal samples included those from cats, mice, and humans. Cat feces were collected from two individuals and stored in plastic zip-top bags. Mouse feces were collected from two individuals by hand using sterile technique and stored in 1.5-mL microcentrifuge tubes. Human feces were collected from five individuals using the Commode Specimen Collection System (Cat#: 02-544-208; Thermo, Carlsbad, CA). Human urine was collected from three male individuals separately into 50-mL centrifuge tubes, and three female individuals separately first into the Commode Specimen Collection System and then transferred to 50-mL centrifuge tubes using sterile technique. Soil samples included soil from the rhizosphere of trees and bare soil. For each type, soil from two sites at the Scripps Coastal Reserve was collected down to a depth of 20 cm using a sterile trowel, and stored in plastic zip-top bags. Water samples included freshwater from two sites at the San Diego River, and seawater from two sites at the Scripps Institution of Oceanography. All water samples were collected and stored in 50-mL centrifuge tubes. Fermented food samples included yogurt and kimchi. For each, two varieties of a single brand were purchased at a local grocery store and transferred to 50-mL centrifuge tubes under sterile conditions. Tissue samples included jejunum tissue from six male mice and six female mice. For each individual, 3.8 cm of the middle small intestine was removed and any particles squeezed out; each tissue section was added to a 2-mL microcentrifuge tube containing 1 mL sterile 1X PBS and 40 mg sterile 1-mm silicone beads, and homogenized at 6,000 rpm for 1 min with a MagNA Lyser (Roche Diagnostics, Santa Clara, CA); the liquid homogenate from intestinal sections from six mice of one gender was pooled to create a single sample. We stored all samples at –80°C within 3 h of collection. To compare limits of detection of microbial cells across kits [29], we included serial dilutions of a mock community containing both bacterial and fungal taxa (*i*.*e*., Zymo Research ZymoBIOMICS Mock Community Standard I, Cat#: D6300). Input cell densities ranged from 140.00-1.40E+09 cells for bacteria, and 2.66-2.66E+07 cells for fungi. Finally, to compare well-to-well contamination [32], we included plasmid-borne, synthetic 16S rRNA gene spike-ins [34] (*i*.*e*., 4 ng of unique spike-in to one well of columns 1-11 in each plate), and at least five extraction blanks per plate.

### DNA extraction

We compared our standardized extraction protocol that uses a 96-sample, magnetic bead cleanup format, the Qiagen MagAttract PowerSoil DNA Isolation Kit (Cat#: 27000-4-KF; Qiagen, Carlsbad, CA), against five other extraction kits: the Qiagen MagAttract PowerSoil Pro DNA Isolation Kit (Cat#: 47109; Qiagen, Carlsbad, CA), the Norgen Stool DNA Isolation Kit (Cat# 65600; Norgen Biotek, Ontario, Canada), the Applied Biosystems MagMAX Microbiome Ultra Nucleic Acid Isolation Kit (Cat#: A42357; Applied Biosystems, Foster City, CA), the Macherey-Nagel NucleoMag Food kit (Cat# 744945.1, Macherey-Nagel, Düren, Germany), and the ZymoBIOMICS 96 MagBead DNA Kit (Cat# D4302, Zymo Research, Irvine, CA). We previously showed that the MagMAX Microbiome kit performs comparably-or-better than our standardized protocol, considering a majority of the criteria included in this benchmark [33]. However, that experiment was focused on establishing the MagMAX kit as an alternative that also allows for downstream RNA-based applications, and did not examine Fungi [33]. Importantly, whereas the PowerSoil, PowerSoil Pro, Norgen, and MagMAX extraction kits employ a 96-deepwell plate format for sample lysis, the NucleoMag Food and Zymo MagBead extraction kits employ a lysis rack, which instead has 12 eight-tube strips arranged in a 96-well rack. This latter format can potentially reduce well-to-well contamination, which is known to occur primarily during the lysis step [32].

For logistical purposes, extractions were performed in two iterations (hereafter referred to as ‘Round 1’ and ‘Round 2’), with a fresh set of samples collected in each instance. Both Round 1 and Round 2 included our standardized protocol as a baseline for comparison. Round 1 centered on the Powersoil Pro and the Norgen kits, and Round 2 the MagMAX, NucleoMag Food, and Zymo MagBead kits. For extraction and following Marotz *et al*. (2017) and Shaffer *et al*. (2021) [28,33], aliquots of each sample were transferred to unique wells of a 96-deepwell extraction plate (or lysis rack). For samples collected with swabs, the entire swab head was broken off into the well (or tube). For liquid samples, we transferred 200 µL. For bulk samples, we used cotton swabs to collect roughly 100 mg of homogenized material and broke the entire swab head off into the well (or tube). For each extraction protocol, all samples, including mock community dilutions, were plated in triplicate. Extractions were performed following the manufacturer’s instructions, with lysis performed using a TissueLyser II (Qiagen, Carlsbad, CA), and bead clean-ups performed using the KingFisher Flex Purification System (ThermoFisher Scientific, Waltham, MA). We stored extracted nucleic acids at –80°C prior to quantification of DNA yield and subsequent sequencing.

### 16S rRNA gene, fungal ITS, and shotgun metagenomics sequencing and data analysis

We prepared DNA for 16S rRNA gene amplicon (16S), fungal ITS gene amplicon (ITS), and shallow shotgun metagenomics sequencing as described previously [10,33,35-38]. Extracts from Round 1 and Round 2 were sequenced separately on distinct runs. For bacterial/archaeal 16S data, raw sequence files were demultiplexed using Qiita [39], sub-operational taxonomic units (sOTUs) generated using Deblur with the default positive filter for 16S data [40], taxonomy assigned using QIIME2’s *feature-classifier* plugin’s *classify-sklearn* method with the pre-fitted classifier trained on GreenGenes V4 (13_8) data [41-44], and phylogeny inferred using QIIME2’s *fragment-insertion* plugin’s *sepp* method with the GreenGenes (13_8) SEPP reference [45-47]. For fungal ITS data, raw sequence files were demultiplexed using Qiita [39], sub-operational taxonomic units (sOTUs) generated using Deblur with a positive filter representing the UNITE8 reference database (dynamic OTUs, including global 97% singletons) [48], and taxonomy assigned using QIIME2’s *feature-classifier* plugin’s *classify-sklearn* method with a classifier trained on the UNITE8 reference database (described above) [41,48]. For shallow shotgun metagenomics data, raw sequence files were demultiplexed using BaseSpace (Illumina, La Jolla, CA), and uploaded to Qiita [39] for additional pre-processing. Demultiplexed sequence data were quality-filtered using fastp [49] and human read depleted by alignment to human reference genome GRCh38 using minimap2 [50]. Filtered reads were aligned to the Web of Life database [51] using bowtie2 [52], and alignment profiles translated to feature-tables using Woltka [53]. Raw sequence data were deposited at the European Nucleotide Archive (accession#: ERP124610) and raw and processed data are available via Qiita (Study ID: 12201). For all three datasets, subsequent normalization of sampling effort and estimation of alpha- and beta-diversity were performed using QIIME2 [9]. Analyses of taxonomic composition and beta-diversity were performed using custom Python scripts. Correlation tests and Kruskal-Wallis tests were performed in R [54]. All processing and analysis code is available on GitHub (github.com/justinshaffer/Extraction_kit_testing).

## Results & Discussion

For each of the five candidate extraction kits tested, we observed similar DNA extraction efficiency to our standardized protocol (hereafter referred to as ‘PowerSoil’), with the exception of the Norgen kit, which had lower yields across all sample types except human milk (Figure 1, Figure S1A). Across the majority of sample types, the PowerSoil Pro, NucleoMag Food, MagMAX Microbiome, and Zymo MagBead kits performed comparably-or-better than PowerSoil (Figure 1, Figure S1A). We also observed similar trends in the number of quality-filtered reads generated from sequencing for each of the five candidate extraction kits as compared to PowerSoil, for 16S, fungal ITS, and shotgun metagenomics data (Figure 2, Figure S1B, Figure S2). Exceptions include the Zymo MagBead kit, which generated fewer high-quality 16S reads across samples from the built environment, water, human urine, and human skin (Figure 2E, Figure S1B), as well as the Norgen kit, which generated fewer high-quality fungal ITS reads across samples from the built environment, food, water, and soil (Figure 2G, Figure S2A). Interestingly, the reduced performance of the Norgen kit in extracting DNA (Figure 1B) did not influence its ability to generate high-quality shotgun metagenomics reads, in-line with PowerSoil (Figure 2L, Figure S2B).

**Figure 1.**
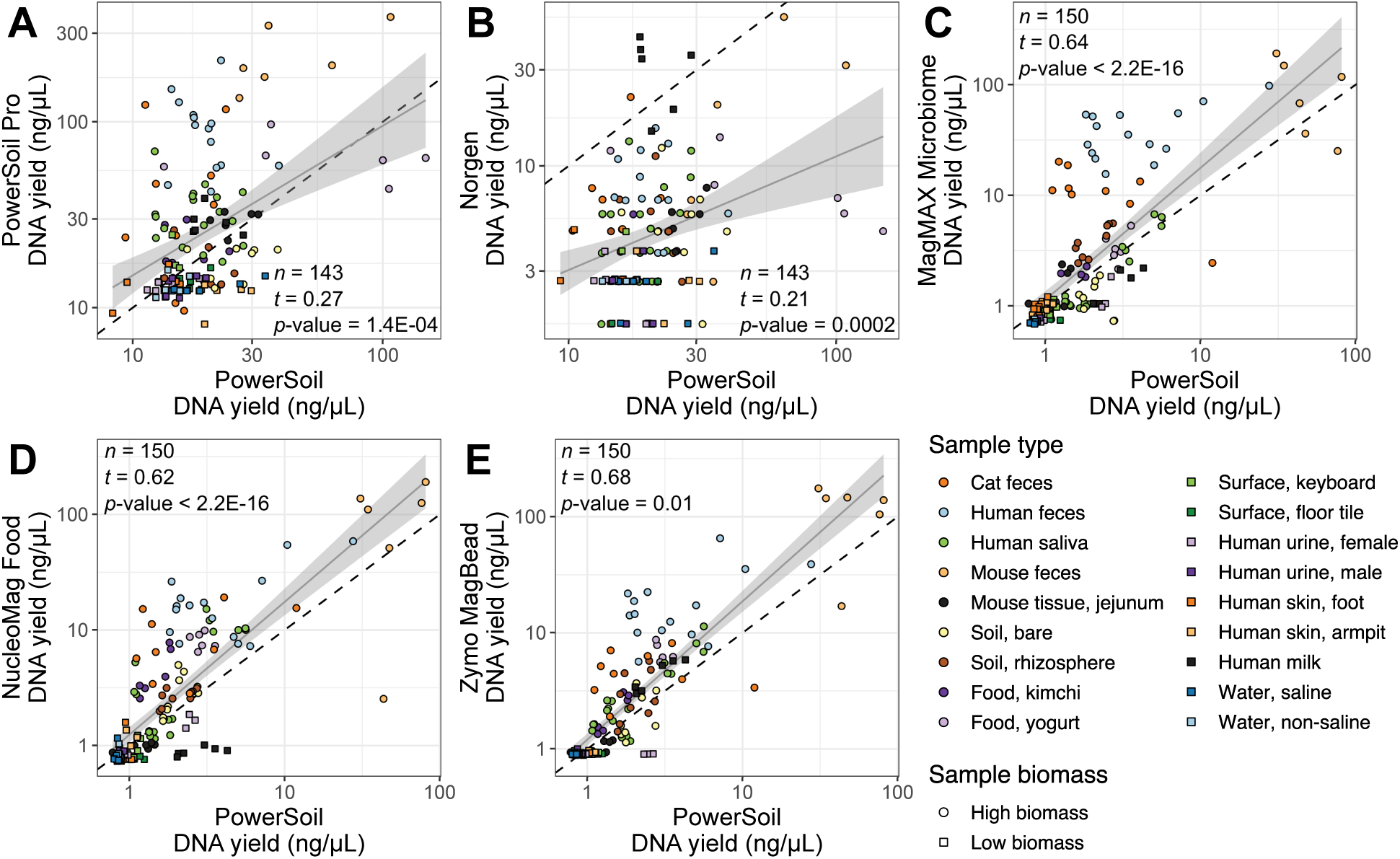
DNA yield (ng/µL) per sample for each candidate extraction kit (*y*-axis) as compared to our standardized protocol (*x*-axis). **(A)** PowerSoil Pro. **(B)** Norgen Stool. **(C)** MagMAX Microbiome. **(D)** NucleoMag Food. **(E)** Zymo MagBead. For all panels, colors indicate sample type and shapes sample biomass, and dotted gray lines indicate 1:1 relationships between methods. Results from tests for correlation between per-sample DNA yield for each respective candidate kit vs. from our standardized protocol, are shown (*t* = Kendall’s tau). For significant correlations, results from a linear model including a 95% confidence interval for predictions are shown. Both axes are presented in a log_10_ scale. A miniaturized, high-throughput Quant-iT PicoGreen dsDNA assay was used for quantification, with a lower limit of 0.1 ng/μL. Yields below this value were estimated by extrapolating from a standard curve.

**Figure 2.**
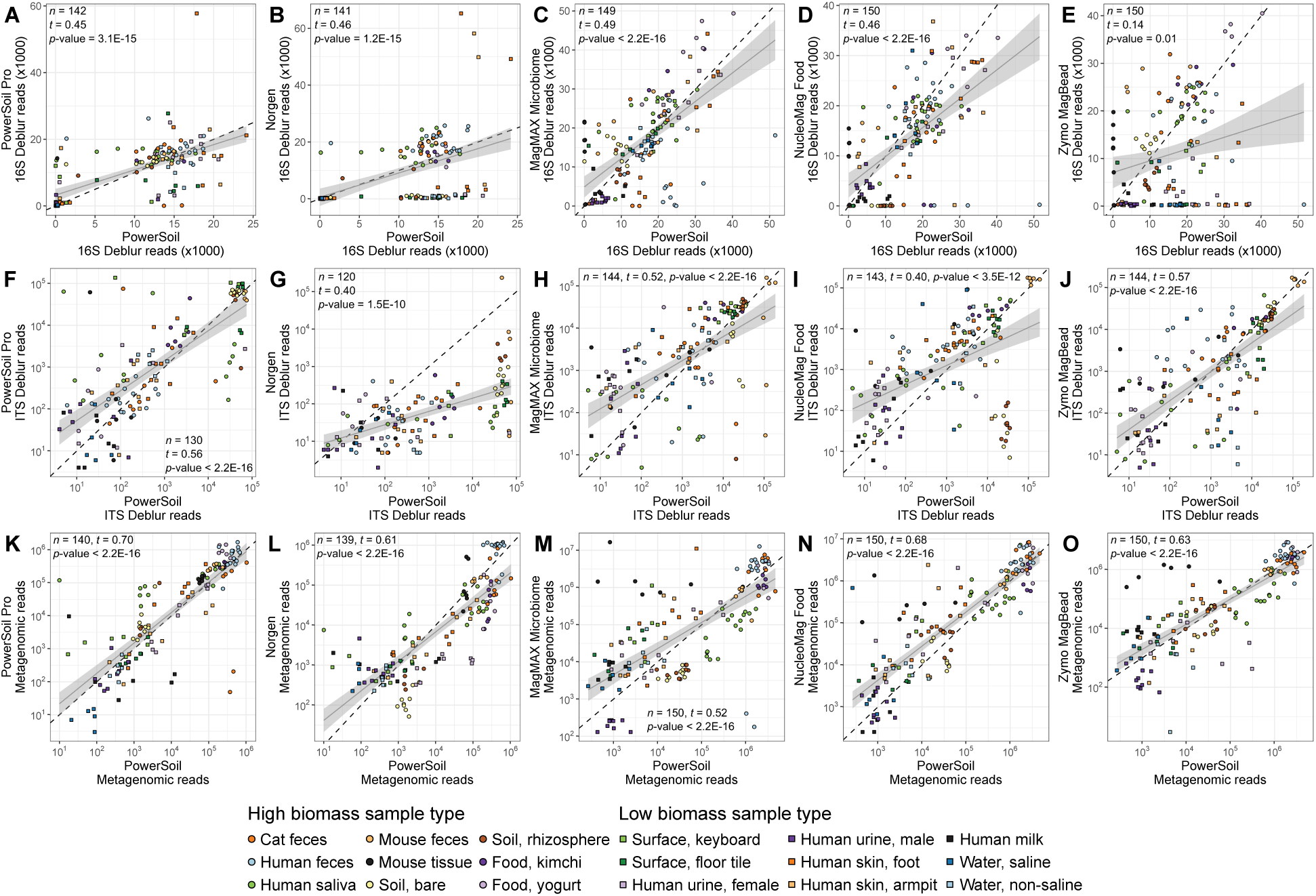
Sequences per sample for each candidate extraction kit (*y*-axis) as compared to our standardized protocol (*x*-axis). **(A-E)** 16S data. **(F-J)** Fungal ITS data. **(K-P)** Shotgun metagenomic data. For each panel, colors indicate sample type and shapes sample biomass, and dotted gray lines indicate a 1:1 relationship between methods. For each dataset, results from tests for correlation between read counts from each respective candidate kit vs. from our standardized protocol, are shown (*t* = Kendall’s tau). For significant correlations, results from a linear model including a 95% confidence interval for predictions are shown. Both axes are presented in a log_10_ scale.

Considering the limit of detection (LOD) of microbial cells, we observed differences in the ability to detect bacteria vs. fungi across the five candidate extraction kits as compared to PowerSoil (Table 1). Compared to PowerSoil, the LOD for bacteria was one order of magnitude lower for the PowerSoil Pro kit, the same for the Zymo MagBead kit, and one order of magnitude higher for MagMAX Microbiome kit (Table 1). However, when considering sample retention following filtering based on LOD thresholds – an important consideration due to the costs of obtaining/processing samples, and for maintaining reasonable sample sizes for analysis – only the PowerSoil Pro and MagMAX Microbiome kits retain ≥80% of samples, similar to PowerSoil (Table 1). The LOD for bacteria for the Norgen and NucleoMag Food kits were much higher, implying that they may not be optimal for profiling of rare taxa (Table 1). The LOD and frequency of sample retention following filtering, for fungi, was similar for all kits as compared to PowerSoil, except for the PowerSoil Pro kit, which had an LOD that was one order of magnitude higher, and also retained only 80% of samples as compared to 100% across all other protocols (Table 1). Surprisingly, the frequency of well-to-well contamination was similar among protocols (Figure 3). This is especially informative considering the unique lysis rack provided by both the NucleoMag Food and Zymo MagBead kits (see Materials & Methods), which we expected to greatly reduce the frequency of well-to-well contamination as compared to lysis in a traditional 96-deepwell plate. We suspect that well-to-well contamination can still occur when using a lysis rack in part due to movement of aerosols during uncapping tubes. We emphasize that without a reduction in well-to-well contamination provided by the lysis rack vs. a traditional plate, the roughly 20-fold increase in processing time to open 96 tubes vs. to unseal a plate (*i*.*e*., ∼100 s vs. 5 s, respectively) argues against adoption of the lysis rack (Figure 3). Future experiments should consider single-tube lysis, which is available for the MagMAX kit (Cat#: A42351), and has been shown to reduce well-to-well contamination [32] although at the cost of increased processing time. Similarly, automated opening/closing of individual, racked tubes, such as that offered by the Matrix Barcoded Storage Tube system (Thermo Scientific, Waltham, MA), should reduce processing time and potential aerosol transfer.

**Table 1.** Limits of detection (LOD) of microbial cells across extraction kits. Titrations of a mock community containing known numbers of cells of bacterial and fungal species (see Materials & Methods) was used to identify the number of per-sample reads needed to meet certain LOD thresholds (*i*.*e*., the percentage of reads mapped to expected taxa vs. contaminants). For each dataset, the read depth corresponding to a threshold of 50% was used for filtering samples prior to community analyses, as recommended [29,33]. The number and percentage of samples retained following filtering based on the read depth for each threshold, as well as LOD estimates for bacterial and fungal cells, are shown for 16S- and ITS data, respectively. Round 1 and Round 2 indicate different sequencing runs, and because sampling effort was not normalized here such to compare absolute read counts, comparisons should not be made across sequencing runs.

| Round | Extraction kit | Threshold (%) | 16S |  |  |  | ITS |  |  |  |
| --- | --- | --- | --- | --- | --- | --- | --- | --- | --- | --- |
|  |  |  | LOD | Read depth | Samples retained | Samples retained (%) | LOD | Read depth | Samples retained | Samples retained (%) |
| 1 | PowerSoil | 50 | 1.40E+03 | 73 | 81 | 86 | 2.66E+00 | 3 | 94 | 100 |
|  |  | 80 | 1.40E+05 | 449 | 55 | 59 | 2.66E+02 | 223 | 45 | 48 |
|  |  | 90 | 1.40E+05 | 1812 | 43 | 46 | > 2.66E+07 | 199,755 | 0 | 0 |
|  |  | 95 | 1.40E+08 | 8676 | 39 | 41 | > 2.66E+07 | 165,759,100,584 | 0 | 0 |
|  | PowerSoil Pro | 50 | 1.40E+02 | 46 | 79 | 84 | 2.66E+01 | 49 | 70 | 80 |
|  |  | 80 | 1.40E+02 | 692 | 61 | 65 | 2.66E+04 | 1,094 | 39 | 45 |
|  |  | 90 | 1.40E+06 | 7520 | 33 | 35 | 2.66E+05 | 18,669 | 14 | 16 |
|  |  | 95 | > 1.40E+09 | 145206 | 0 | 0 | > 2.66E+07 | 698,158 | 0 | 0 |
|  | Norgen Stool | 50 | 1.40E+07 | 1960 | 18 | 19 | 2.66E+00 | 2 | 88 | 100 |
|  |  | 80 | 1.40E+07 | 4705 | 12 | 13 | 2.66E+05 | 850 | 7 | 8 |
|  |  | 90 | 1.40E+08 | 8224 | 10 | 10 | > 2.66E+07 | 314,049,798 | 0 | 0 |
|  |  | 95 | 1.40E+08 | 14215 | 9 | 10 | > 2.66E+07 | > 1.00E+12 | 0 | 0 |
| 2 | PowerSoil | 50 | 1.40E+02 | 88 | 93 | 97 | 2.66E+00 | 3 | 91 | 100 |
|  |  | 80 | 1.40E+02 | 3224 | 80 | 83 | 2.66E+00 | 6 | 91 | 100 |
|  |  | 90 | > 1.40E+09 | 88894 | 0 | 0 | 2.66E+00 | 12 | 86 | 95 |
|  |  | 95 | > 1.40E+09 | 6228705 | 0 | 0 | 2.66E+00 | 24 | 75 | 82 |
|  | MagMAX Microbiome | 50 | 1.40E+03 | 1286 | 79 | 82 | 2.66E+00 | 5 | 92 | 100 |
|  |  | 80 | 1.40E+04 | 4147 | 66 | 69 | 2.66E+00 | 479 | 64 | 70 |
|  |  | 90 | 1.40E+04 | 8970 | 63 | 66 | > 2.66E+07 | 451,442 | 0 | 0 |
|  |  | 95 | 1.40E+04 | 19454 | 39 | 41 | > 2.66E+07 | 176,467,642,943 | 0 | 0 |
|  | NucleoMag Food | 50 | 1.40E+05 | 8379 | 61 | 64 | 2.66E+00 | 1 | 88 | 100 |
|  |  | 80 | 1.40E+05 | 14121 | 45 | 47 | 2.66E+00 | 2 | 88 | 100 |
|  |  | 90 | 1.40E+06 | 19430 | 27 | 28 | 2.66E+02 | 74 | 69 | 78 |
|  |  | 95 | 1.40E+07 | 26321 | 18 | 19 | > 2.66E+07 | 26,901,301 | 0 | 0 |
|  | Zymo MagBead | 50 | 1.40E+02 | 489 | 42 | 44 | 2.66E+00 | 1 | 90 | 100 |
|  |  | 80 | 1.40E+02 | 1838 | 32 | 33 | 2.66E+00 | 4 | 90 | 100 |
|  |  | 90 | 1.40E+06 | 4531 | 22 | 23 | 2.66E+01 | 466 | 38 | 42 |
|  |  | 95 | 1.40E+06 | 11445 | 10 | 10 | > 2.66E+07 | 5,179,906,841 | 0 | 0 |

**Figure 3.**
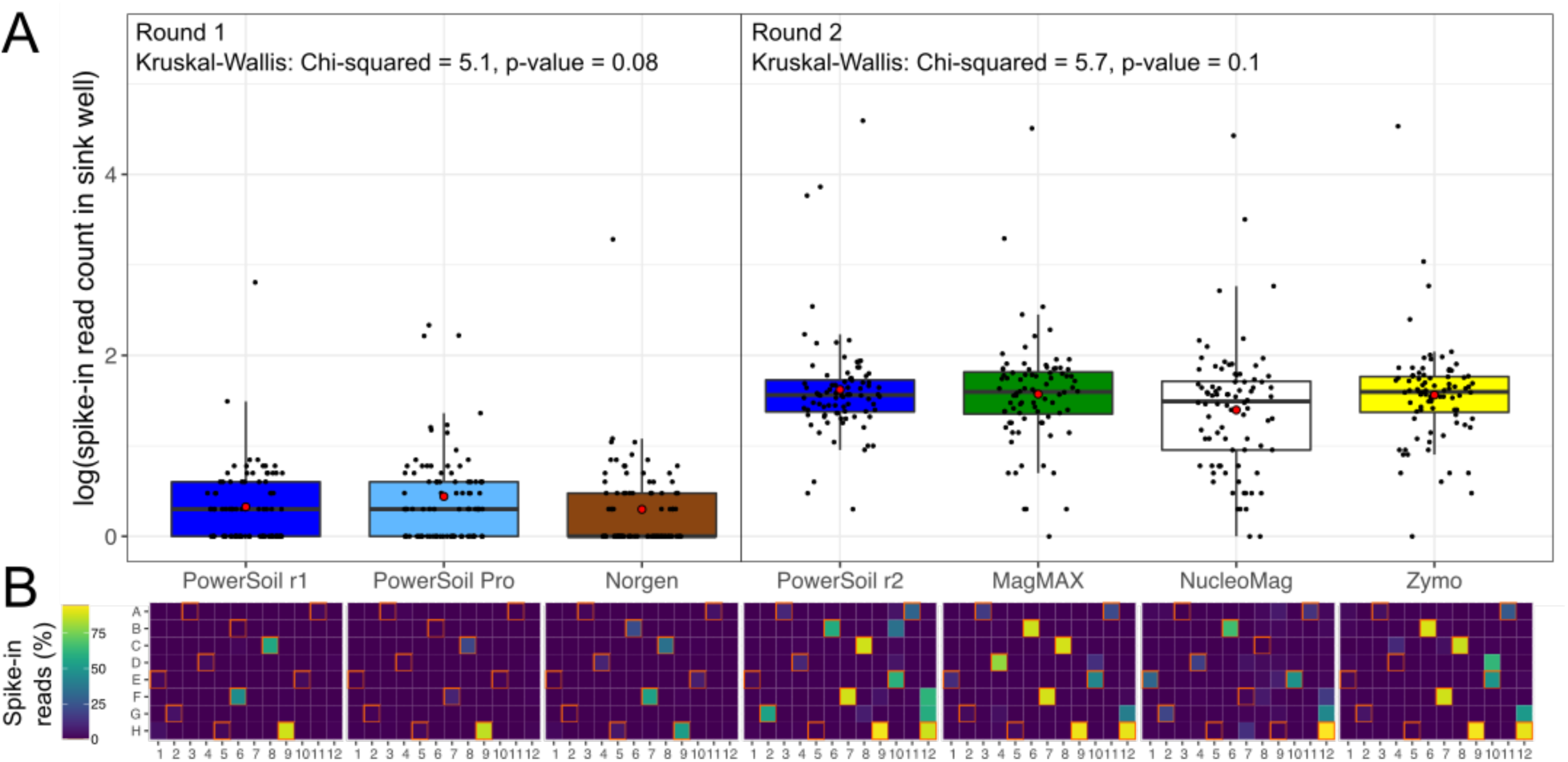
Well-to-well contamination across candidate extraction kits as compared to our standardized protocol. Plasmids harboring synthetic 16S sequences were spiked into a single well per plate column (*i*.*e*., 1-11) of each high-biomass sample plate prior to extraction. **(A)** The number of reads matching synthetic 16S sequences was quantified for all wells that did not receive a spike-in (*i*.*e*., sink wells). Round 1 and Round 2 indicate different sequencing runs, and because sampling effort was not normalized here such to compare absolute read counts, comparisons should not be made across sequencing runs. For each sequencing run, results from a Kruskal-Wallis test are shown. **(B)** The percentage of spike-in reads among all reads per well shown as a heatmap. Wells into which plasmids were spiked (*i*.*e*., source wells) are outlined in orange.

Considering the taxonomic composition of microbial communities across samples, we observed a greater degree of taxon bias among extraction kits as compared to PowerSoil for fungal taxa (*i*.*e*., ITS data) as compared to bacterial/archaeal taxa (*i*.*e*., 16S and shotgun metagenomics data) (Figure 4C, Figure 4D). This is likely due in part to the relatively diverse morphologies among fungal spores and propagules compared to those of bacteria/archaea, which may be variably compromised among distinct lysis approaches. Both the PowerSoil Pro and MagMAX Microbiome kits recovered the greatest number of exclusive fungal genera (*i*.*e*., those not recovered by other protocols), with each taxon set representing roughly 19% of all fungal genera recovered in a given round of extractions (Figure 4C, Figure 4D). Similarly, both the PowerSoil Pro and MagMAX Microbiome kits shared a greater number of exclusive fungal genera with PowerSoil, as compared to the other candidate extraction kits (Figure 4C, Figure 4D). We observed a similar trend in our 16S data, except that for both rounds of extraction PowerSoil recovered the greatest number of exclusive bacterial/archaeal genera as compared to any candidate kit (Figure 4A, Figure 4B). For our shotgun metagenomics data, all candidate extraction kits except for the Norgen kit recovered a greater number of exclusive bacterial/archaeal species than PowerSoil (Figure 4E, Figure 4F). The PowerSoil Pro and NucleoMag Food kits recovered the greatest percentage of exclusive species, with the MagMAX Microbiome and Zymo MagBead kits recovering only slightly less (Figure 4E, Figure 4F). The PowerSoil Pro kit shared the greatest percentage of exclusive species with PowerSoil (16%), whereas the NucleoMag Food and MagMAX Microbiome kits shared much less (∼1%) (Figure 4F).

**Figure 4.**
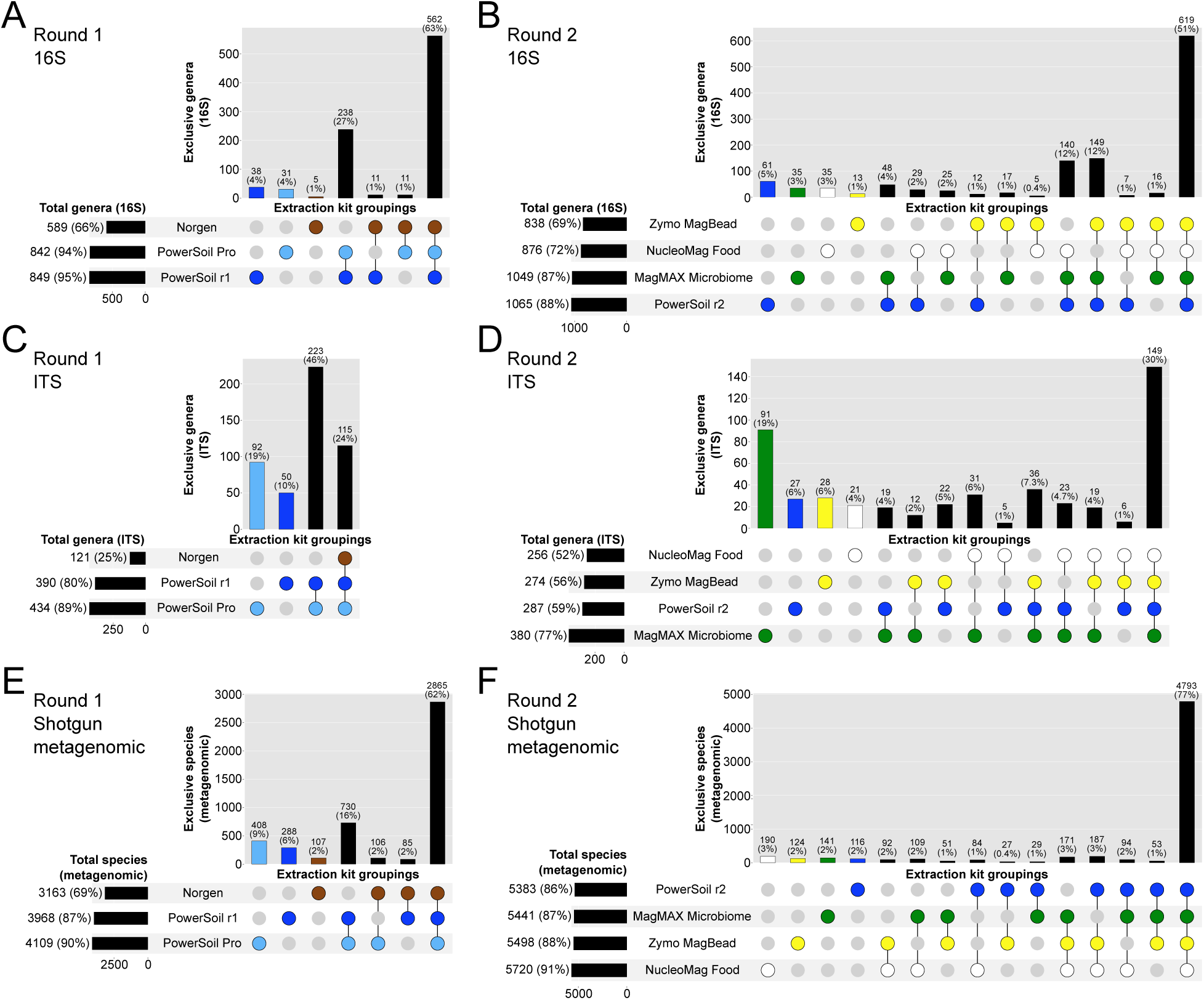
Taxonomic bias among extraction protocols. Upset plots showing **(A, B)** genera for bacterial/archaeal 16S data, **(C, D)** genera for fungal ITS data, and **(E, F)** species for bacterial/archaeal metagenomics data, highlighting taxa shared among extraction protocols. Values indicate counts and percentages are respective to all taxa across all protocols. Associations representing <5 taxa were excluded for clarity. Round 1 and Round 2 indicate different sequencing runs, and because sampling effort was not normalized here such to compare absolute taxon counts, comparisons of counts (*i*.*e*., vs. percentages) should not be made across sequencing runs.

For non-exclusive taxa, we observed strong correlations in the relative abundance estimates from each candidate extraction kit as compared to PowerSoil (Figure 5). For 16S data, the strongest correlation was observed between PowerSoil and the MagMAX Microbiome kit (Kendall’s tau = 0.67) (Figure 5C), followed by the Zymo MagBead kit (tau = 0.66) (Figure 5E), and the Norgen kit (tau = 0.64) (Figure 5B). For fungal ITS data, the strongest correlation was observed with the Zymo MagBead kit (tau = 0.58) (Figure 5J), followed by the MagMAX Microbiome kit (tau = 0.47) (Figure 4H), and the PowerSoil Pro kit (tau = 0.46) (Figure 5F). For shotgun metagenomics data, the strongest correlation was observed with the Zymo MagBead kit (tau = 0.68) (Figure 5O), followed by the NucleoMag Food kit (Kendall’s tau = 0.67) (Figure 5N), and the MagMAX Microbiome kit (tau = 0.59) (Figure 5M).

**Figure 5.**
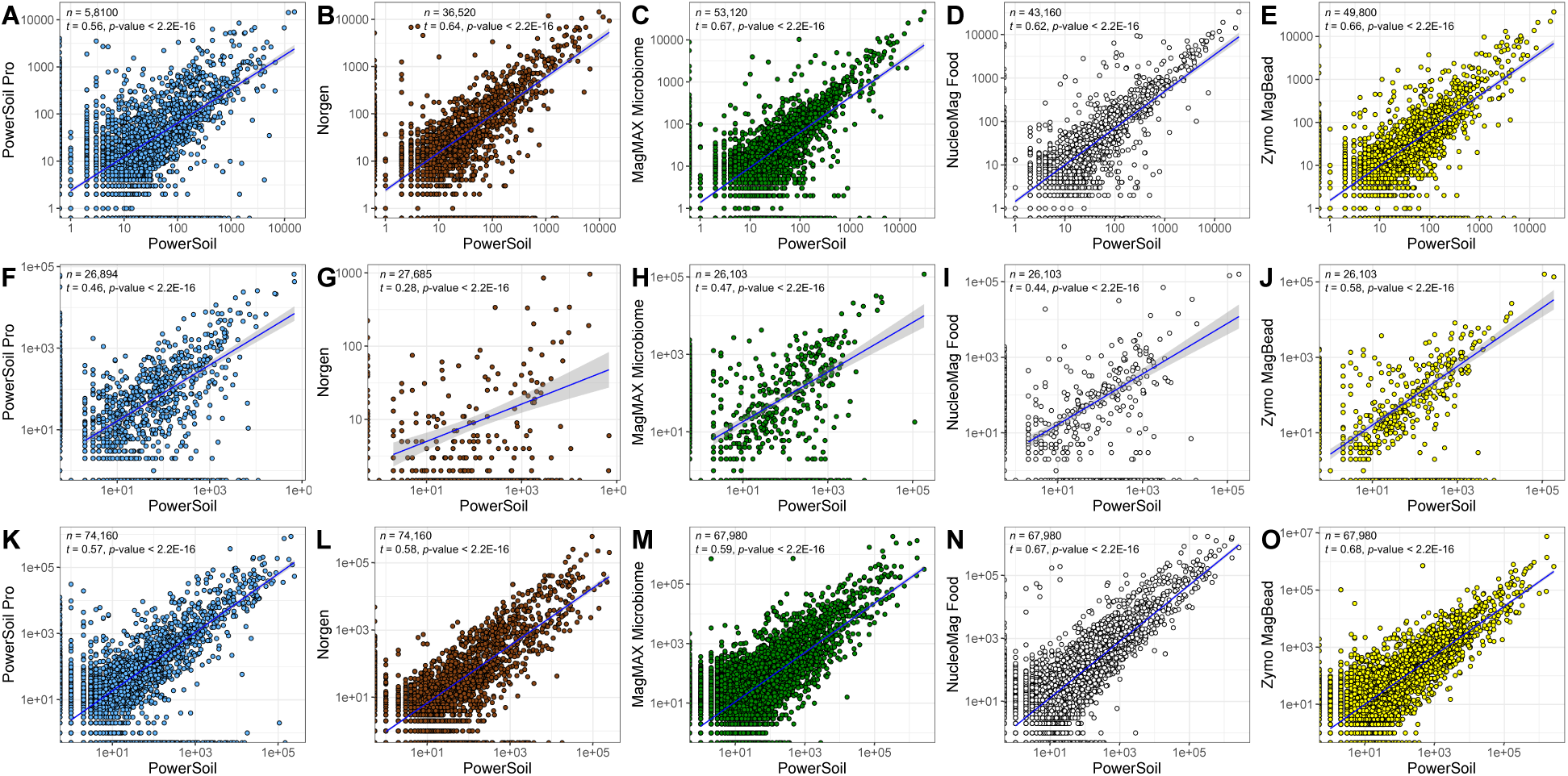
Taxon relative abundances across all samples for each candidate extraction kit (*y-*axis) as compared to our standardized protocol (*x*-axis). Colors indicate candidate kits. **(A-E)** 16S data, where each point represents a bacterial/archaeal genus. **(F-J)** Fungal ITS data, where each point represents a fungal genus. **(K-O)** Shotgun metagenomic data, where each point represents a bacterial/archaeal genus. For each dataset, results from tests for correlation between taxon abundances from each respective candidate kit vs. our standardized protocol, are shown (*t* = Kendall’s tau). For significant correlations, results from a linear model including a 95% confidence interval for predictions are shown. Both axes are presented in a log_10_ scale.

We also observed strong correlations in estimates of microbial community alpha-diversity from each candidate extraction kit as compared to PowerSoil (Figure 6A-E), and note in general correlations were stronger for 16S and shotgun metagenomics data as compared to for ITS data (Figure 6F-J). Specifically, correlations between candidate kits and PowerSoil for 16S alpha-diversity (*i*.*e*., Faith’s Phylogenetic Diversity [PD]) were all strong (tau >0.75), except for the Norgen kit, which had the weakest correlation and also the greatest sample dropout from normalization (Figure 6B). Similarly, correlations between candidate kits and PowerSoil for fungal ITS alpha-diversity (*i*.*e*., Fisher’s alpha) were also strong (tau >0.60), except for the PowerSoil Pro kit, which had a relatively weak relationship (Figure 6F), and the Norgen kit, which had no relationship and also significant sample dropout (Figure 6G). Correlations between candidate kits and PowerSoil for shotgun metagenomics alpha-diversity (*i*.*e*., Faith’s PD) were all strong (tau >0.65), and sample dropout was minimal for all protocols (Figure 6K-O).

**Figure 6.**
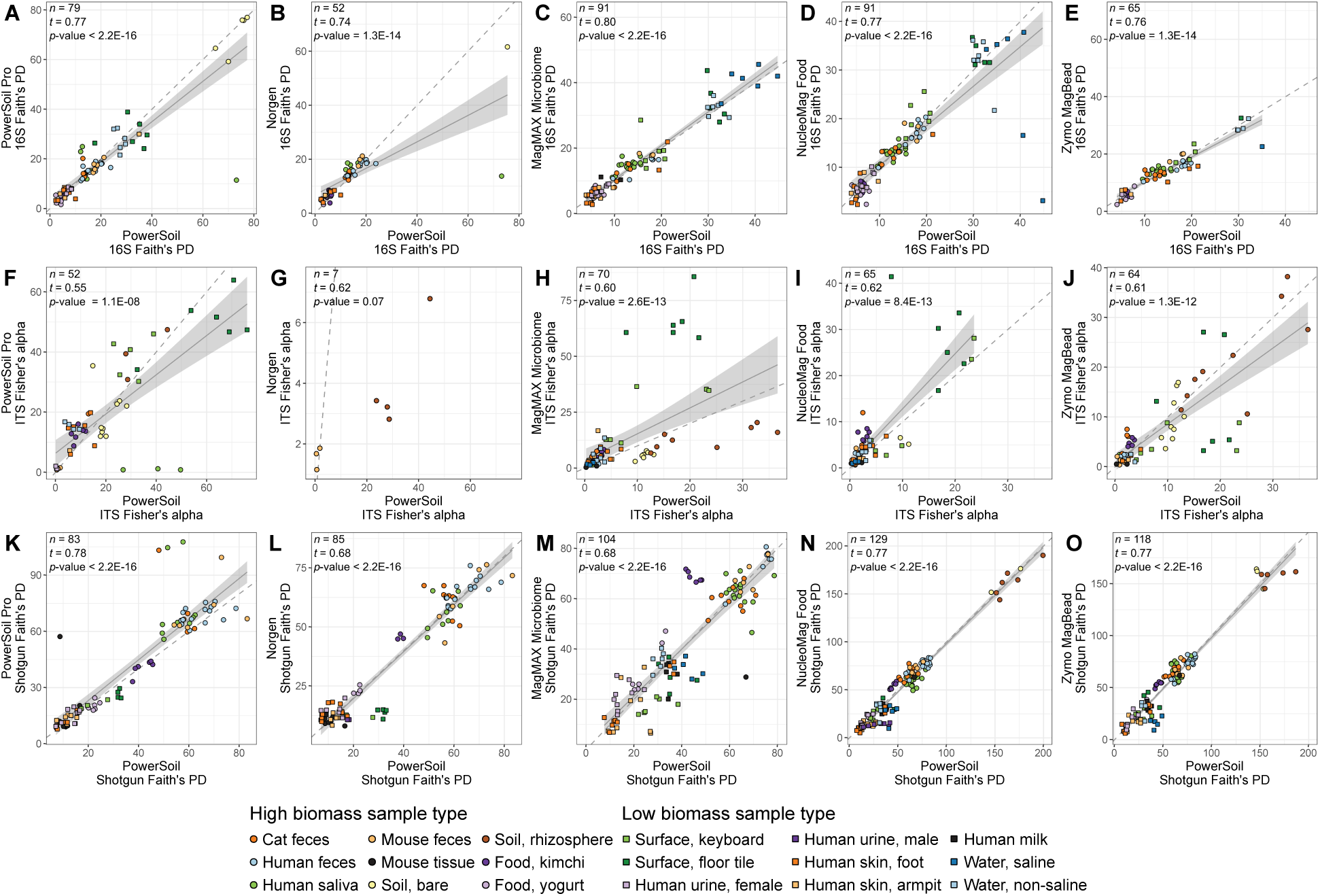
Alpha-diversity per sample across sample types for each candidate extraction kit (*y*-axis) as compared to our standardized protocol (*x*-axis). **(A-E)** 16S data. **(F-J)** Fungal ITS data. **(K-P)** Shotgun metagenomic data. For each panel, colors indicate sample type and shapes sample biomass, and dotted gray lines indicate a 1:1 relationship between methods. Results from tests for correlation between alpha-diversity values from each respective candidate kit vs. from our standardized protocol, are shown (*t* = Kendall’s tau). For significant correlations, results from a linear model including a 95% confidence interval for predictions are shown. Sample types absent from any panel lacked representation by the respective candidate extraction kit and our standardized protocol.

With respect to microbial community composition, we found variation explained by bias among extraction protocols to be negligible compared with that explained by host subject identity (*i*.*e*., 1-2 orders of magnitude weaker in explaining beta-diversity) (Table 2). For 16S and shotgun metagenomics data, the variation explained by extraction protocol is one order of magnitude weaker for presence/absence metrics vs. abundance-based metrics (Table 2). This supports our analysis of variation among technical replicates from the same sample, which we observed to be small for all extraction protocols across 16S (Figure S3), fungal ITS (Figure S4), and shotgun metagenomics data (Figure S5). We also observed strong correlations in microbial community beta-diversity (*i*.*e*., sample-sample distances) from each candidate extraction kit as compared to PowerSoil, and note that as for alpha-diversity, in general correlations were stronger for 16S and shotgun metagenomics data as compared to ITS data (Table S1, Table S2, Table S3). Specifically, correlations in sample-sample distances between each of the five candidate kits and PowerSoil for 16S data were strong (rho >0.75), except for in low-biomass samples processed with the Norgen kit, which exhibited no relationship with PowerSoil for two-of-four distance metrics examined (Table S1). For ITS data, correlations in sample-sample distances were consistently weaker for low biomass samples vs. high biomass samples, for all candidate extraction kits and distance metrics examined. The PowerSoil Pro kit alone had correlation coefficients >0.50 for low biomass samples and >0.75 for high biomass samples (Table S2). For shotgun metagenomics data, whereas correlations in sample-sample distances were also consistently weaker for low biomass samples vs. high biomass samples, the magnitude of the difference was smaller as compared to ITS data, and correlations for high biomass samples were strong (rho >0.85) for all candidate extraction kits and distance metrics examined (Table S3).

**Table 2.** <u>Assessment of</u>factors influencing microbial community beta-diversity in this study. Results are from forward, stepwise model selection, following Shaffer *et al*. (2021) [33]. Values are based on permutation tests of variation explained by redundancy analysis (*n* = 5,000 runs), done separately for unique distance metrics for 16S, ITS, and shotgun metagenomic data. The full model included extraction round (*i*.*e*., round 1 vs. 2), sample biomass (*i*.*e*., high vs low biomass), sample type, host subject identity, and extraction kit as model variables. 16S data were rarefied to 10,000 quality-filtered reads per sample or had samples with fewer than 10,000 reads excluded when using RPCA distances (*n* = 640 samples). Fungal ITS data were rarefied to 630 quality-filtered reads per sample or had samples with fewer than 630 reads excluded when using RPCA distances (*n* = 978 samples). Shotgun metagenomic data were rarefied to 2,100 host- and quality-filtered reads per sample or had samples with fewer than 2,100 reads excluded when using RPCA distances (*n* = 1,044 samples). Rarefaction depths were selected to maintain at least 75% samples (50% for fungal ITS data) from both high- and low-biomass datasets. AIC: Akaike information criterion; df: degrees of freedom; RPCA: Robust principal components analysis (*i*.*e*., Robust Aitchison distance).

| Data type | Distance metric | Factor | Adjusted R<sup>2</sup> | df | AIC | F | p-value |
| --- | --- | --- | --- | --- | --- | --- | --- |
| 16S | Unweighted UniFrac | Host identity | 0.93 | 60 | -979.83 | 152.03 | 0.0002 |
|  | Unweighted UniFrac | Extraction kit | 0.001 | 6 | -988.95 | 3.20 | 0.0004 |
|  | Weighted UniFrac | Host identity | 0.84 | 60 | -428.10 | 58.62 | 0.0002 |
|  | Weighted UniFrac | Extraction kit | 0.02 | 6 | -533.75 | 19.27 | 0.0002 |
|  | Jaccard | Host identity | 0.96 | 60 | -1234.51 | 231.05 | 0.0002 |
|  | Jaccard | Extraction kit | 0.002 | 6 | -1254.66 | 4.92 | 0.0002 |
|  | RPCA | Host identity | 0.89 | 60 | -644.89 | 86.15 | 0.0002 |
|  | RPCA | Extraction kit | 0.004 | 6 | -663.81 | 4.73 | 0.0002 |
| ITS | Jaccard | Host identity | 0.80 | 59 | -191.58 | 33.90 | 0.0002 |
|  | Jaccard | Extraction kit | 0.02 | 6 | -235.28 | 8.51 | 0.0002 |
|  | RPCA | Host identity | 0.72 | 59 | -30.05 | 22.32 | 0.0002 |
|  | RPCA | Extraction kit | 0.02 | 6 | -62.07 | 6.64 | 0.0002 |
| Metagenomic | Unweighted UniFrac | Host identity | 0.92 | 65 | -1063.58 | 141.03 | 0.0002 |
|  | Unweighted UniFrac | Extraction kit | 0.005 | 6 | -1107.59 | 8.79 | 0.0002 |
|  | Weighted UniFrac | Host identity | 0.87 | 65 | -672.15 | 81.26 | 0.0002 |
|  | Weighted UniFrac | Extraction kit | 0.02 | 6 | -807.08 | 24.46 | 0.0002 |
|  | Jaccard | Host identity | 0.93 | 65 | -1195.00 | 168.79 | 0.0002 |
|  | Jaccard | Extraction kit | 0.003 | 6 | -1225.27 | 6.57 | 0.0002 |
|  | RPCA | Host identity | 0.85 | 65 | -579.25 | 70.95 | 0.0002 |
|  | RPCA | Extraction kit | 0.01 | 6 | -633.10 | 10.40 | 0.0002 |

Importantly, whereas agreement with PowerSoil regarding the results of analyzing 16S or shotgun metagenomics data is desirable, deviation from PowerSoil among the five candidate extraction kits based on fungal ITS data was expected. In that regard, the MagMAX Microbiome kit alone consistently maintains a high degree of correlation with PowerSoil for both 16S and shotgun metagenomics data (Figure 2C, Figure 2M, Figure 5C, Figure 5M, Figure 6C, Figure 6M), while also maintaining a relatively high number of samples from both high- and low biomass subsets following normalization, across all three data layers (Table S1, Table S2, Table S3). The NucleoMag Food and Zymo MagBead kits have slightly stronger correlations with PowerSoil as compared to MagMAX for shotgun metagenomics data for certain analyses of microbial community diversity (Figure 6M-O, Table S1, Table S3). However, the MagMAX Microbiome kit recovered the greatest number of exclusive fungal genera while also sharing the greatest number of exclusive fungal genera with PowerSoil (Figure 4D). Although the PowerSoil Pro kit also recovered a similar frequency of exclusive fungal genera (Figure 4C), the increased sample dropout following normalization for that kit vs. the MagMAX Microbiome kit (*i*.*e*., particularly for low biomass samples) (Table S1, Table S2, Table S3), increased processing time (*i*.*e*., approximately 3.5 h for PowerSoil and PowerSoil Pro vs. 1.0 h for MagMAX Microbiome), as well as the increased cost of consumables (*i*.*e*., 3.5X reagent reservoirs and 7X tips for PowerSoil and PowerSoil Pro vs. MagMAX Microbiome) combined with previous work showing that the MagMAX kit can also extract high-quality RNA from similar samples [33], argue strongly for the use of the MagMAX Microbiome kit. However, the PowerSoil Pro kit is a good alternative if there are no downstream RNA applications, and if time and cost are not important factors.

## Conclusions

We conclude that the MagMAX Microbiome extraction kit is comparable to our standardized PowerSoil protocol with respect to characterizing microbial community composition using both 16S and shotgun metagenomic data, and is optimal compared to other candidate kits and our standardized protocol for doing so using fungal ITS data, as it recovers the greatest number of unique fungal genera. The PowerSoil Pro kit is a good alternative, as it excels in the same regards, but it does not extract RNA and is a more time- and cost-intensive protocol, as compared to the MagMAX Microbiome kit. Regardless, data from the PowerSoil, PowerSoil Pro, and MagMAX Microbiome extraction kits should allow for comparisons such as meta-analysis across 16S, ITS, and shotgun metagenomics data produced using those protocols and downstream processing and analytical methods similar to those used here. In addition to recovering a greater number of fungal taxa, the more rapid processing time, and use of fewer consumables highlight the MagMAX Microbiome kit as a comparable and efficient alternative to the PowerSoil protocol that also optimizes downstream applications, including fungi.

### Future perspective

Future efforts should continue to focus on optimizing microbiological and molecular methods that capture all organisms in a sample regardless of their evolutionary history, and from a diversity of sample types such as those examined here. Such methods provide invaluable resources and should serve as gold standards to be adopted widely by the community [56]. In parallel, further advances in computational methods should focus on reducing technical effects in meta-analyses across studies using distinct methods [6,39]. In concert, such advances will allow us to maximize our understanding of microbial communities and to harness that knowledge in part to foster human and environmental sustainability.

### Ethical conduct of research

The human subjects work conducted here is approved through UCSD IRB#150275.

### Executive summary

1. We compared our previously established, standardized protocol for DNA extraction against five alternative DNA extraction kits.
2. We included a diverse panel of sample types ranging from host-associated to environmental.
3. We also included controls for detecting well-to-well contamination and the limit of detection of microbial cells.
4. We observed sample type-specific differences in DNA extraction efficiency among all extraction protocols.
5. Sample type and host identity were stronger drivers of microbial community beta-diversity as compared to the extraction protocol used.
6. We identify one protocol that generates high-quality fungal ITS data, and produces 16S- and shotgun metagenomics data with high similarity to our established protocol with respect to microbial community alpha-diversity, beta-diversity, and taxonomic composition.
7. The similarity between the optimal protocol and our existing one will allow for meta-analyses across both with negligible technical bias.

