## Supplementary Material for "A comparison of six DNA extraction protocols for 16S, ITS, and shotgun metagenomic sequencing of microbial communities"

**Supplementary tables and figures.**

**Table S1.** Mantel correlations in sample-sample distances between each candidate extraction kit and our standardized protocol, for bacterial/archaeal 16S sequence data. Data were rarefied to the maximum read depth that maintained 75% of samples, or had samples with fewer than that number of reads excluded when using RPCA distances (*i.e.*, high biomass samples: 12,690 reads; low biomass samples: 3,295 reads).

| Data type | Sample biomass | Extraction kit | *n* | Distance metric | Pearsons's *r* | *p*-value |
|---|---|---|---|---|---|---|
| 16S | High biomass | PowerSoil Pro | 45 | Unweighted UniFrac | 0.79 | 0.0002 |
| | | | | Weighted UniFrac | 0.77 | 0.0002 |
| | | | | Jaccard | 0.84 | 0.0002 |
| | | | | RPCA | 0.84 | 0.0002 |
| | | Norgen | 42 | Unweighted UniFrac | 0.87 | 0.0002 |
| | | | | Weighted UniFrac | 0.86 | 0.0002 |
| | | | | Jaccard | 0.92 | 0.0002 |
| | | | | RPCA | 0.86 | 0.0002 |
| | | MagMAX | 40 | Unweighted UniFrac | 0.90 | 0.0002 |
| | | | | Weighted UniFrac | 0.93 | 0.0002 |
| | | | | Jaccard | 0.95 | 0.0002 |
| | | | | RPCA | 0.90 | 0.0002 |
| | | NucleoMag | 45 | Unweighted UniFrac | 0.90 | 0.0002 |
| | | | | Weighted UniFrac | 0.94 | 0.0002 |
| | | | | Jaccard | 0.96 | 0.0002 |
| | | | | RPCA | 0.93 | 0.0002 |
| | | Zymo | 45 | Unweighted UniFrac | 0.91 | 0.0002 |
| | | | | Weighted UniFrac | 0.90 | 0.0002 |
| | | | | Jaccard | 0.96 | 0.0002 |
| | | | | RPCA | 0.96 | 0.0002 |
| | Low biomass | PowerSoil Pro | 28 | Unweighted UniFrac | 0.90 | 0.0002 |
| | | | | Weighted UniFrac | 0.91 | 0.0002 |
| | | | | Jaccard | 0.91 | 0.0002 |
| | | | | RPCA | 0.88 | 0.0002 |
| | | Norgen | 7 | Unweighted UniFrac | 0.33 | 0.2 |
| | | | | Weighted UniFrac | 0.85 | 0.008 |
| | | | | Jaccard | 0.81 | 0.002 |
| | | | | RPCA | 0.07 | 0.8 |
| | | MagMAX | 48 | Unweighted UniFrac | 0.92 | 0.0002 |
| | | | | Weighted UniFrac | 0.93 | 0.0002 |
| | | | | Jaccard | 0.91 | 0.0002 |
| | | | | RPCA | 0.83 | 0.0002 |
| | | NucleoMag | 44 | Unweighted UniFrac | 0.90 | 0.0002 |
| | | | | Weighted UniFrac | 0.90 | 0.0002 |
| | | | | Jaccard | 0.89 | 0.0002 |
| | | | | RPCA | 0.82 | 0.0002 |
| | | Zymo | 18 | Unweighted UniFrac | 0.89 | 0.0002 |
| | | | | Weighted UniFrac | 0.87 | 0.0002 |
| | | | | Jaccard | 0.89 | 0.0002 |
| | | | | RPCA | 0.81 | 0.0002 |

**Table S2.** Mantel correlations in sample-sample distances between each candidate extraction kit and our standardized protocol, for fungal ITS sequence data. Data were rarefied to the maximum read depth that maintained 50% of samples, or had samples with fewer than that number of reads excluded when using RPCA distances (*i.e.*, high biomass samples: 1,491 reads; low biomass samples: 344 reads).

| Data type | Sample biomass | Extraction kit | *n* | Distance metric | Pearsons's *r* | *p*-value |
|---|---|---|---|---|---|---|
| ITS | High biomass | PowerSoil Pro | 25 | Jaccard | 0.80 | 0.0002 |
| | | | | RPCA | 0.77 | 0.0002 |
| | | Norgen | 6 | Jaccard | 0.68 | 0.02 |
| | | | | RPCA | 0.59 | 0.06 |
| | | MagMAX | 36 | Jaccard | 0.70 | 0.0002 |
| | | | | RPCA | 0.71 | 0.0002 |
| | | NucleoMag | 33 | Jaccard | 0.58 | 0.0002 |
| | | | | RPCA | 0.64 | 0.0002 |
| | | Zymo | 45 | Jaccard | 0.74 | 0.0002 |
| | | | | RPCA | 0.73 | 0.0002 |
| | Low biomass | PowerSoil Pro | 26 | Jaccard | 0.67 | 0.0002 |
| | | | | RPCA | 0.65 | 0.0004 |
| | | MagMAX | 33 | Jaccard | 0.49 | 0.0002 |
| | | | | RPCA | 0.31 | 0.003 |
| | | NucleoMag | 32 | Jaccard | 0.50 | 0.0002 |
| | | | | RPCA | 0.24 | 0.02 |
| | | Zymo | 19 | Jaccard | 0.23 | 0.02 |
| | | | | RPCA | 0.18 | 0.1 |

**Table S3.** Mantel correlations in sample-sample distances between each candidate extraction kit and our standardized protocol, for bacterial/archaeal shotgun metagenomic sequence data. Data were rarefied to the maximum read depth that maintained 75% of samples, or had samples with fewer than that number of reads excluded when using RPCA distances (*i.e.*, high biomass samples: 38,000 reads; low biomass samples: 600 reads).

| Data type | Sample biomass | Extraction kit | *n* | Distance metric | Pearsons's *r* | *p*-value |
|---|---|---|---|---|---|---|
| Metagenomic | High biomass | PowerSoil Pro | 57 | Unweighted UniFrac | 0.95 | 0.0002 |
| | | | | Weighted UniFrac | 0.92 | 0.0002 |
| | | | | Jaccard | 0.95 | 0.0002 |
| | | | | RPCA | 0.88 | 0.0002 |
| | | Norgen | 42 | Unweighted UniFrac | 0.99 | 0.0002 |
| | | | | Weighted UniFrac | 0.94 | 0.0002 |
| | | | | Jaccard | 0.99 | 0.0002 |
| | | | | RPCA | 0.93 | 0.0002 |
| | | MagMAX | 54 | Unweighted UniFrac | 0.97 | 0.0002 |
| | | | | Weighted UniFrac | 0.87 | 0.0002 |
| | | | | Jaccard | 0.97 | 0.0002 |
| | | | | RPCA | 0.94 | 0.0002 |
| | | NucleoMag | 64 | Unweighted UniFrac | 0.98 | 0.0002 |
| | | | | Weighted UniFrac | 0.94 | 0.0002 |
| | | | | Jaccard | 0.99 | 0.0002 |
| | | | | RPCA | 0.94 | 0.0002 |
| | | Zymo | 59 | Unweighted UniFrac | 0.98 | 0.0002 |
| | | | | Weighted UniFrac | 0.90 | 0.0002 |
| | | | | Jaccard | 0.99 | 0.0002 |
| | | | | RPCA | 0.95 | 0.0002 |
| | Low biomass | PowerSoil Pro | 25 | Unweighted UniFrac | 0.80 | 0.0002 |
| | | | | Weighted UniFrac | 0.86 | 0.0002 |
| | | | | Jaccard | 0.87 | 0.0002 |
| | | | | RPCA | 0.70 | 0.0002 |
| | | Norgen | 30 | Unweighted UniFrac | 0.60 | 0.0002 |
| | | | | Weighted UniFrac | 0.46 | 0.0002 |
| | | | | Jaccard | 0.51 | 0.0002 |
| | | | | RPCA | 0.56 | 0.0002 |
| | | MagMAX | 53 | Unweighted UniFrac | 0.72 | 0.0002 |
| | | | | Weighted UniFrac | 0.78 | 0.0002 |
| | | | | Jaccard | 0.80 | 0.0002 |
| | | | | RPCA | 0.73 | 0.0002 |
| | | NucleoMag | 55 | Unweighted UniFrac | 0.69 | 0.0002 |
| | | | | Weighted UniFrac | 0.87 | 0.0002 |
| | | | | Jaccard | 0.76 | 0.0002 |
| | | | | RPCA | 0.67 | 0.0002 |
| | | Zymo | 49 | Unweighted UniFrac | 0.61 | 0.0002 |
| | | | | Weighted UniFrac | 0.83 | 0.0002 |
| | | | | Jaccard | 0.79 | 0.0002 |
| | | | | RPCA | 0.35 | 0.003 |

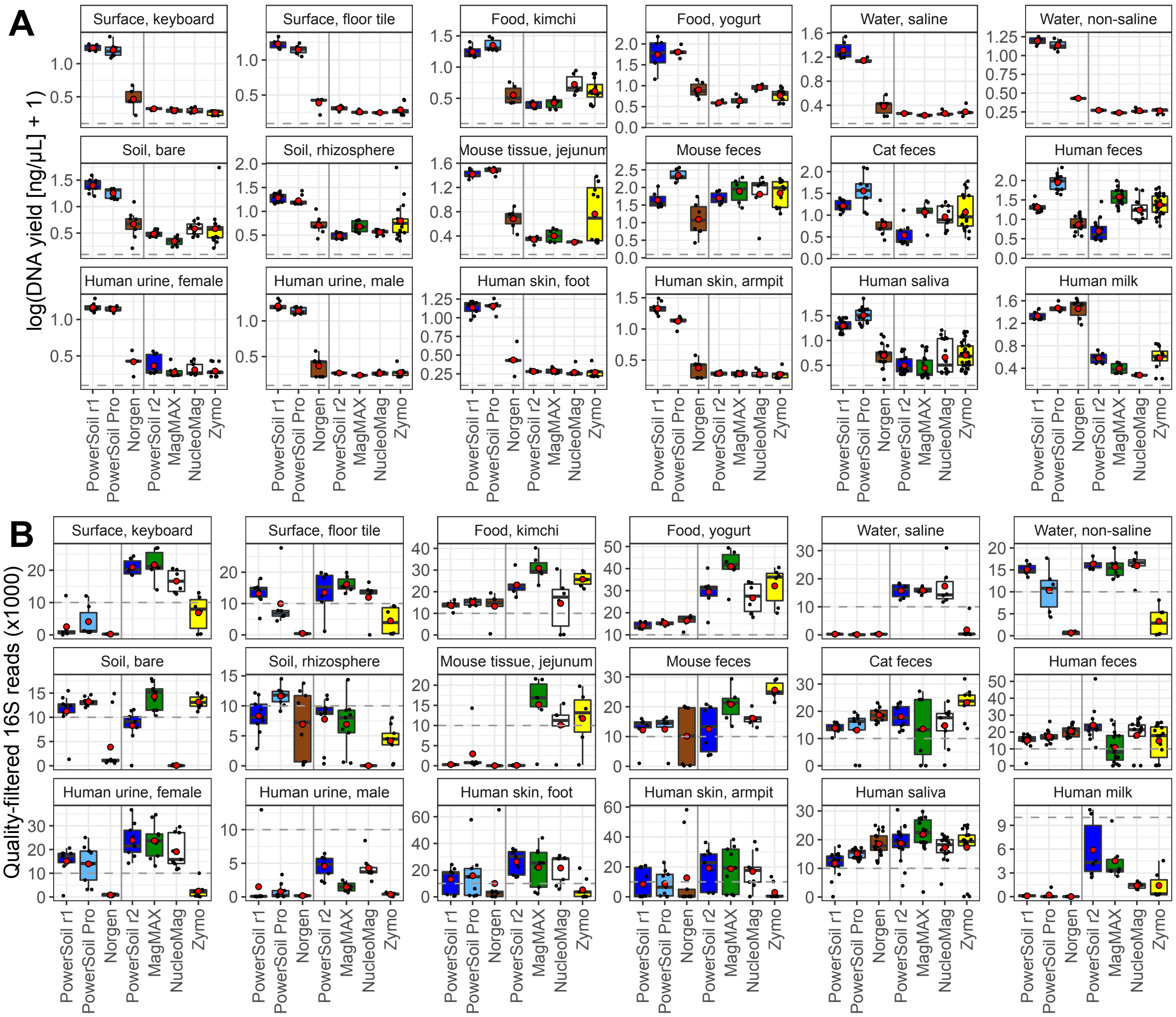


**Figure S1. (A)** Average concentration of DNA (ng/µL) across extraction protocols for each sample type (*n* = 1,184 samples). Red circles indicate group means. A miniaturized, high-throughput Quant-iT PicoGreen dsDNA assay was used, with a lower limit of 0.1 ng/µL indicated by the horizontal, dotted gray line in each panel. Yields below this value were estimated by extrapolating from a standard curve. **(B)** Average number of quality-filtered sequences for 16S data (*n* = 1,039 samples). Dashed lines indicate our expectation of 10,000 reads from human fecal samples. For both panels, red circles indicate means, and vertical gray lines separate different sequencing runs. As sampling effort was not normalized here, such to maintain absolute values, comparisons should not be made across sequencing runs.

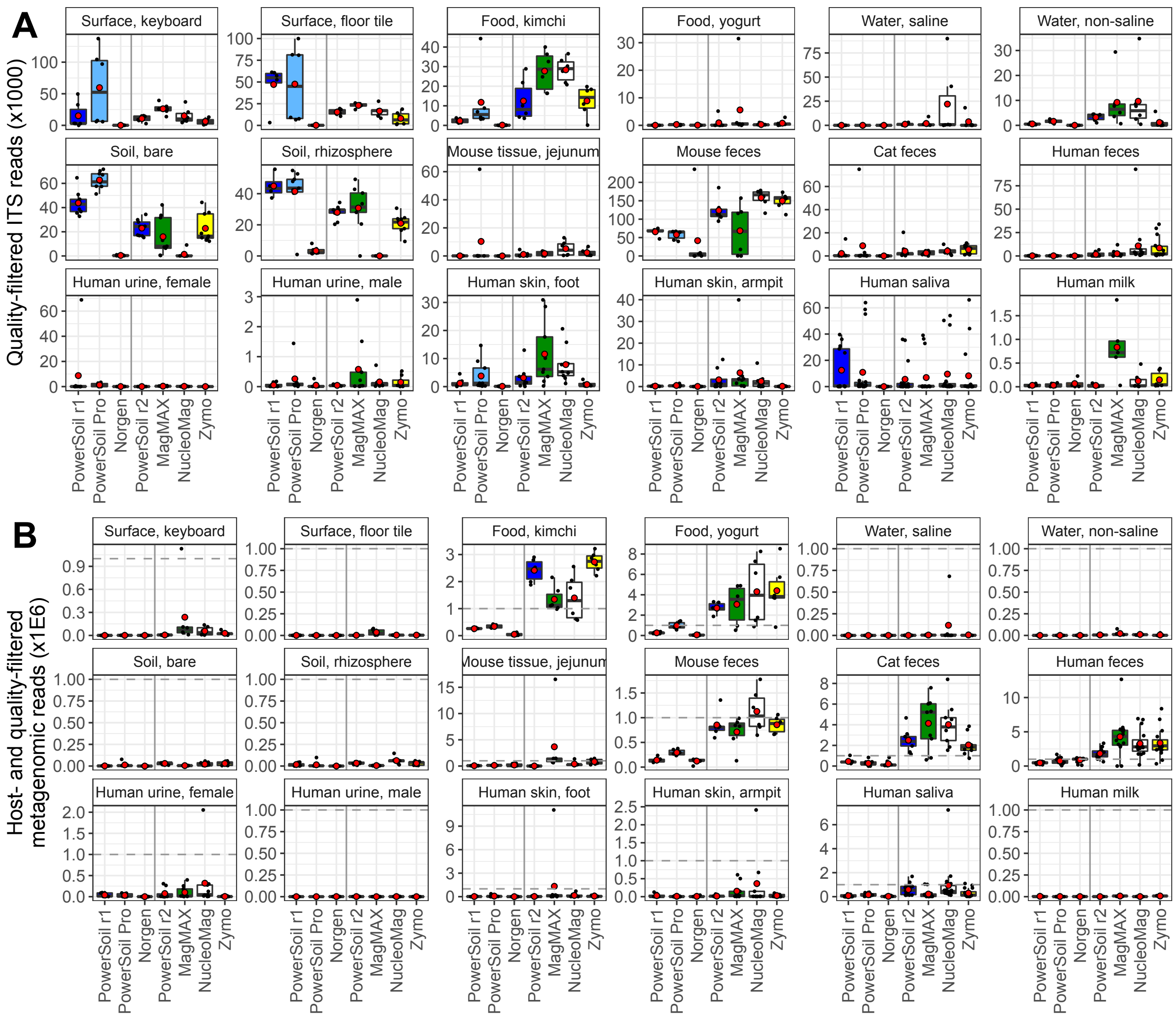

**Figure S2.** Sequences per sample across extraction protocols and sample types. **(A)** Average number of quality-filtered sequences for fungal ITS data (*n* = 991 samples). **(B)** Average number of host- and quality-filtered sequences for bacterial/archaeal metagenomic data (*n* = 1,037 samples). Dashed lines indicate our expectation of 1,000,000 reads from human fecal samples. For both panels, red circles indicate means, and vertical gray lines separate different sequencing runs. As sampling effort was not normalized here, such to maintain absolute read counts, comparisons should not be made across sequencing runs.

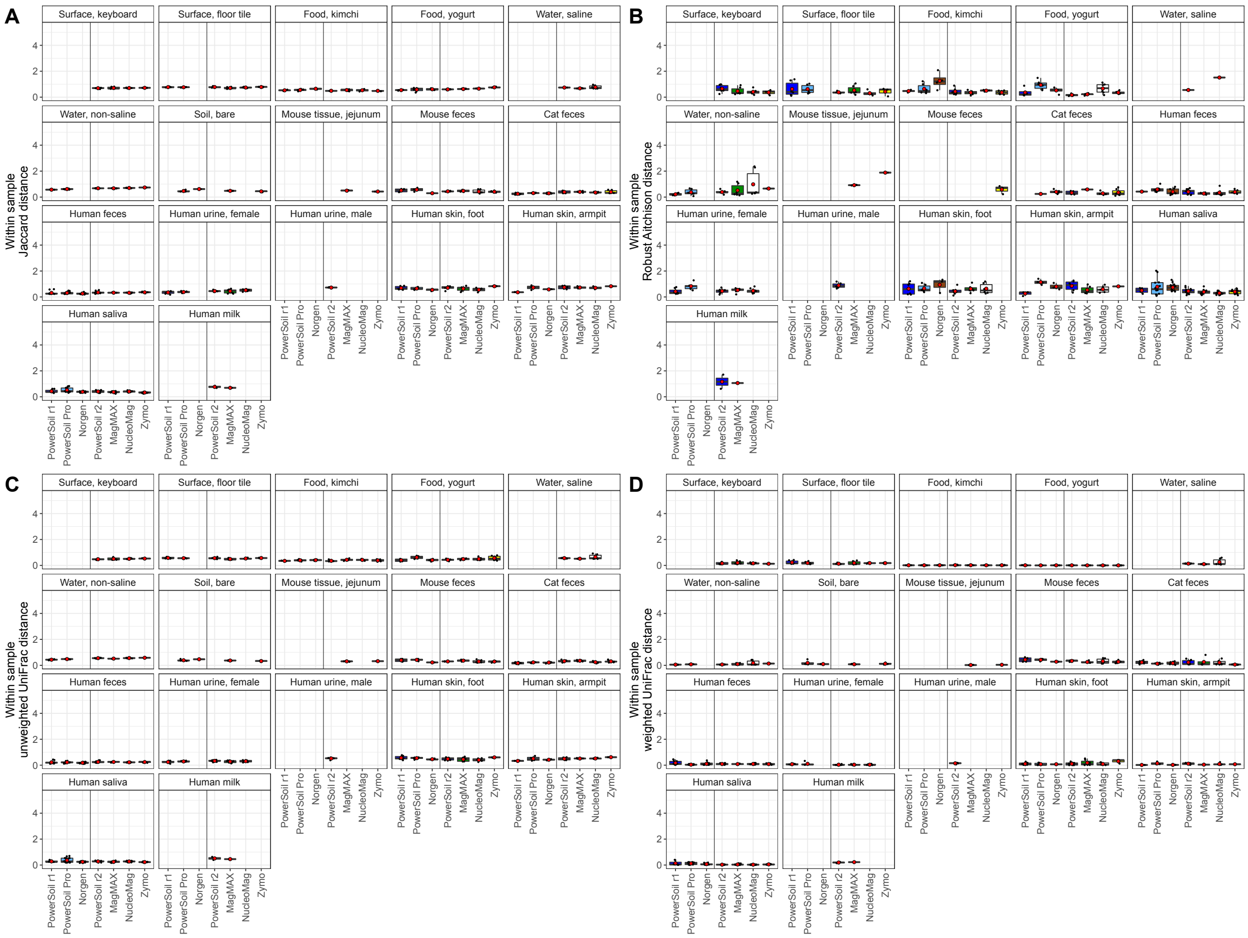


**Figure S3.** Within-sample variation across extraction kits, for bacterial/archaeal 16S data. Microbial community beta-diversity among replicate extractions of the same source sample was estimated using (A) Jaccard distance, (B) RPCA distance, (C) unweighted UniFrac distance, and (D) weighted UniFrac distance. Data were rarefied to the maximum read depth that maintained 75% of samples, or had samples with fewer than that number of reads excluded when using RPCA distances (*i.e.*, high biomass samples: 12,690 reads; low biomass samples: 3,295 reads).

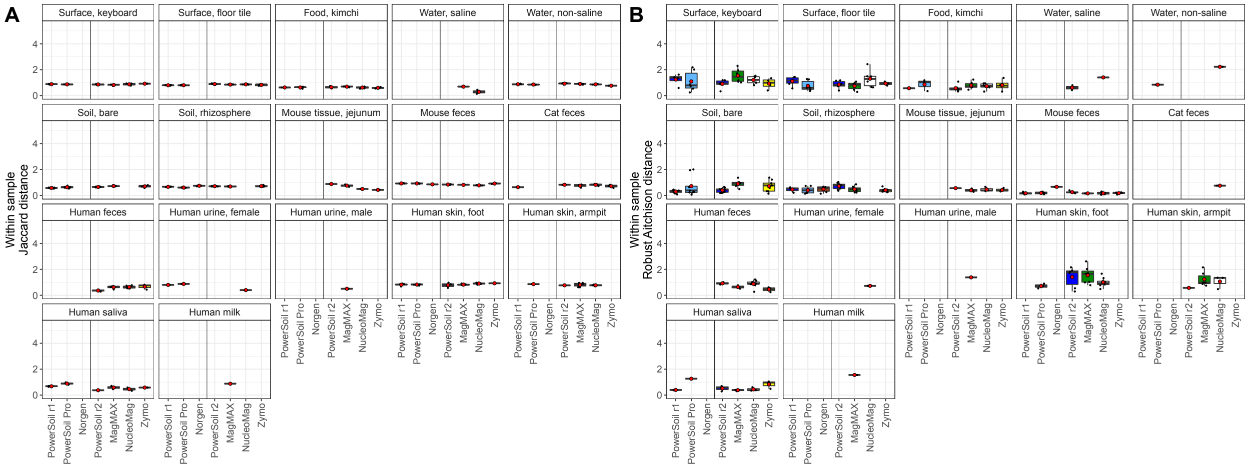


**Figure S4.** Within-sample variation across extraction kits, for fungal ITS data. Fungal community beta-diversity among replicate extractions of the same source sample was estimated using (A) Jaccard distance, and (B) RPCA distance. Data were rarefied to the maximum read depth that maintained 50% of samples, or had samples with fewer than that number of reads excluded when using RPCA distances (*i.e.*, high biomass samples: 1,491 reads; low biomass samples: 344 reads).

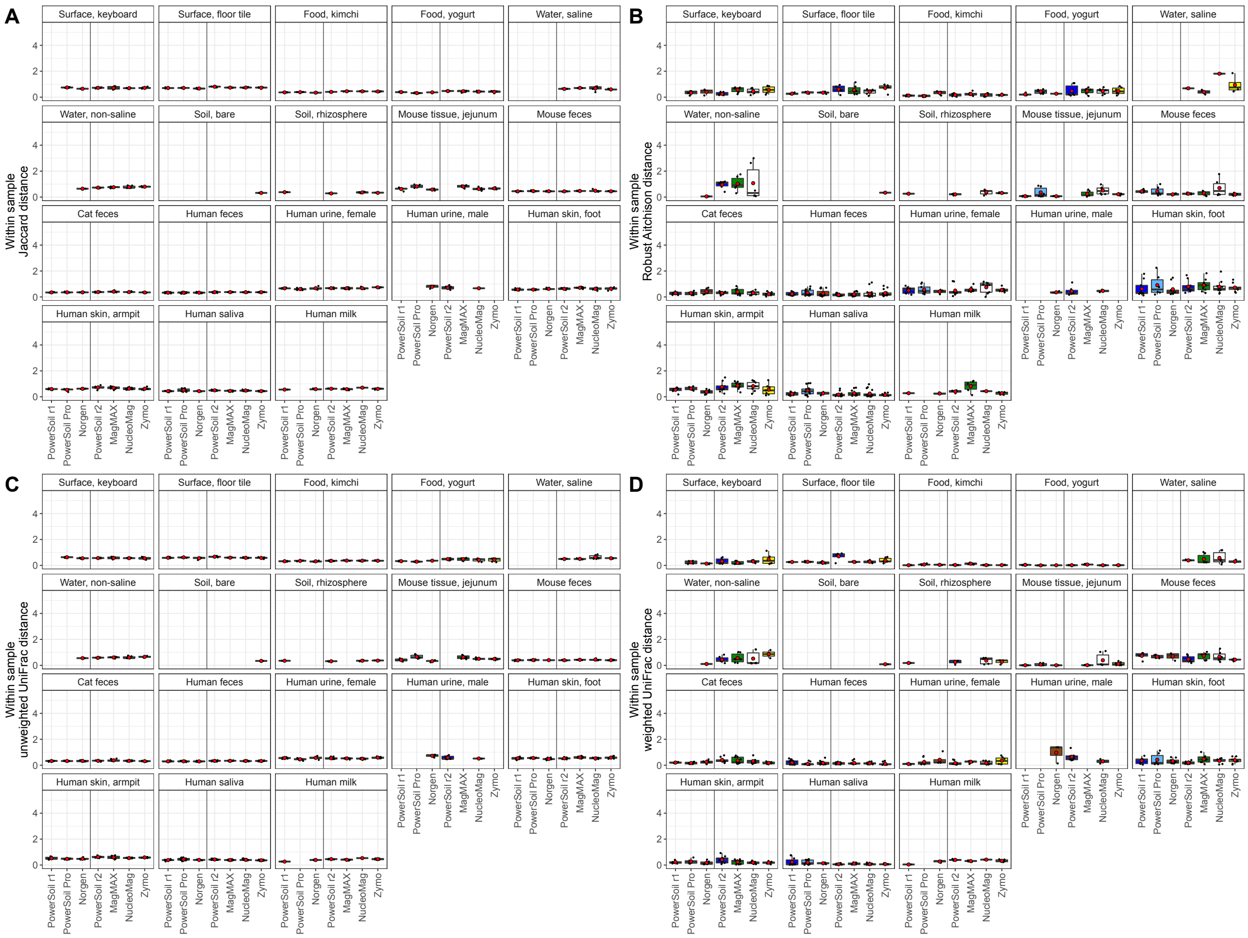


**Figure S5.** Within-sample variation across extraction kits, for bacterial/archaeal shotgun metagenomic sequence data. Microbial community beta-diversity among replicate extractions of the same source sample was estimated using (A) Jaccard distance, (B) RPCA distance, (C) unweighted UniFrac distance, and (D) weighted UniFrac distance. Data were rarefied to the maximum read depth that maintained 75% of samples, or had samples with fewer than that number of reads excluded when using RPCA distances (*i.e.*, high biomass samples: 38,000 reads; low biomass samples: 600 reads).
